# NAD^+^ controls circadian rhythmicity during cardiac aging

**DOI:** 10.1101/2023.11.02.565150

**Authors:** Bryce J. Carpenter, Margaux Lecacheur, Yannick N. Mangold, Kai Cui, Stefan Günther, Pieterjan Dierickx

## Abstract

Disruption of the circadian clock as well as reduced NAD^+^ levels are both hallmarks of aging. While circadian rhythms and NAD^+^ metabolism have been linked in heart disease, their relationship during cardiac aging is less clear. Here, we show that aging leads to disruption of diurnal gene expression in the heart. Long-term supplementation with the NAD^+^ precursor nicotinamide riboside (NR) boosts NAD^+^ levels, reprograms the diurnal transcriptome and reverses naturally occurring cardiac enlargement in aged female mice. In addition, complete abolishment of NAD^+^ levels in CMs impairs PER2::luc oscillations, which is rescued by NR supplementation. These findings reveal an essential role for NAD^+^ in regulation of the cardiac circadian clock upon aging, which opens up new avenues to counteract age-related cardiac disorders.

## Introduction

The circadian clock is an endogenous time-keeping system that allows for the anticipation of rhythmic stimuli such as light/dark and feeding/fasting cycles and is observed in nearly all mammals^1^. The underlying molecular machinery consists of a transcriptional/translational feedback loop including transcription factors such as BMAL1, CLOCK, REV-ERBα/β, and PER2 amongst others. The clock drives rhythmic mRNA expression of tissue-specific genes both *in vivo* and *in vitro*. These oscillations can persist in the absence of external clues but are responsive to changes in the environment, such as light, nutrients and food. In the heart specifically, around 6% of the transcriptome is rhythmically expressed^2^, and many clock output genes are implicated in metabolic pathways. Indeed, circadian clocks are tightly linked to metabolic homeostasis, which is best illustrated by genetic disruption of the clock leading to obesity and metabolic syndrome, as well as cardiovascular defects in rodents^3,4^. In humans, environmental clock disruption (via, for example, shift-work or jet-travel) is also correlated with the occurrence of a higher risk for metabolic and cardiovascular disorders^3^.

Age is one of the greatest risk factors for both likelihood and severity of heart disease, pairing with common pathological parameters such as hypertrophy, reduced cardiac output, and increased oxidative stress^5^. As a highly metabolically active organ, the heart is susceptible to alterations in fuel selection and processing, incidence of which also increases with age, like diabetes and obesity^6^. Interestingly, the heart is known to swap to a more fetal-like metabolic state during cardiac failure, increasing reliance on glycolysis^7,8^. Aging has also been linked to dampening of clock gene expression and circadian processes^9–11^, which would further correlate to deregulated metabolism and heart disorders^12,13^. Indeed, whole body KO of *Bmal1* in mice leads to premature aging^14^ and we have previously shown that KO of both *Rev-erbs* in the heart (cardiomyocyte-specific RevDKO) induces metabolic dysregulation and dilated cardiomyopathy, leading to early lethality^4^. The prominent cofactor NAD^+^ has been shown to fall under circadian control in the heart^4,15^, and also decreases with age across many species and tissues^11–16^. Generally, these alterations in NAD^+^ levels are largely mediated by either redox status or enzymes that consume NAD^+^, such as sirtuins, poly ADP-ribose polymerases (PARPs), and CD38^16^. In addition, NAD^+^ feeds back into the clock via modulating activity of SIRT1, an NAD^+^-dependent deacetylase working in tandem with the BMAL1:CLOCK dimer that forms the positive arm of the core clock loop^17^. This includes regulation of *Nampt* expression, which encodes for the major rate-limiting enzyme in the salvage of nicotinamide (NAM) into NAD^+^, critical to heart NAD^+^ levels as the heart is incapable of *de novo* synthesis^18^. Alternatively, cardiac NAD^+^ can be generated from nicotinamide riboside (NR) by NMRK2, which is upregulated in human and murine heart failure^19^ while NAMPT decreases^18^. A mechanistic study in the murine liver has shown that during aging, lower SIRT1 activity causes higher PER2 stability and prolonged repression of BMAL1, thereby reducing rhythmicity of clock output genes and demonstrating at least one link between dampened oscillations and NAD^+^ loss over age^20^. Since high amplitude oscillations are linked to health^21^, increasing the amplitude of the core clock and its output might be a viable strategy to reverse aging.

Boosting NAD^+^ levels via supplementation with NAD^+^ precursors has proven beneficial in a number of age-associated diseases including murine heart failure models^22–26^. However, whether and how NAD^+^ boosting affects circadian rhythmicity in the heart is not known. In this study, we examine the effect of modulation of NAD^+^ levels to investigate whether increasing NAD^+^ levels can improve clock function in the heart with age. *In vitro*, we demonstrate alterations in cardiac PER2::luc protein oscillations as a function of age using isolated adult cardiomyocytes. By using orally-supplemented NR in aging female mice, a treatment strategy we have previously shown to improve cardiac function^19^, we identified that transcriptional oscillations of cardiac clock output genes normally affected during aging can be reprogrammed. Additionally, we verify *in vitro* that manipulation of NAD^+^, with boosting via NR treatment or loss by pharmacological inhibition of NAMPT, is able to severely impact PER2::luc oscillations in neonatal cardiomyocytes. These results reveal that NAD^+^ plays an essential role in regulating the diurnal cardiac transcriptome. Moreover, NR supplementation reduced age-related cardiac enlargement and stress. Together, this data suggests a link between NAD^+^ metabolism, the circadian clock, and cardiac health during aging, which could be harnessed to ameliorate cardiac defects often occurring in aged individuals.

## Results and discussion

Aging has been reported to affect circadian gene expression in different tissues^9–11^. To test its effect on diurnal gene expression in the heart, we harvested hearts from young (8 weeks) and old (1 year) female mice at *Zeitgeber* (ZT)22 (night) and ZT10 (day), two timepoints at which the core clock factors *Bmal1* and *Clock* are highly and lowly expressed, respectively, as determined by cardiac qRT-PCR on an independent full circadian female young cohort (**Fig. 1a and Supplemental Fig. 1)**. We assessed gene expression via bulk mRNA-seq and observed 1,231 diurnally expressed genes in young mice and 302 in old ones (**Fig.1b and Supplemental Table 1**, *Adj. p=0*.*05*). Of those, 142 genes oscillated in both, while 1,089 and 160 were unique diurnal in young and old mice respectively (**Fig. 1b-c**). This shows that, in line with other organs^11^, fewer genes oscillate in the hearts of older animals. Young-specific diurnal genes were enriched for pathways such as ECM-receptor interaction, focal adhesion, PI3K-Akt signaling, while oscillators unique to old hearts were enriched for pathways in cancer, Ras signaling, and MAPK signaling (**Fig. 1d**). Genes that displayed diurnal expression in both young and old hearts were enriched for the Circadian Rhythm pathway only (**Fig. 1d**). Indeed, core clock gene expression was only mildly affected in aged animals (**Fig. 1e**). These results suggest that rhythmic expression of clock output genes is both reduced and rewired in the aging heart.

**Fig. 1.**
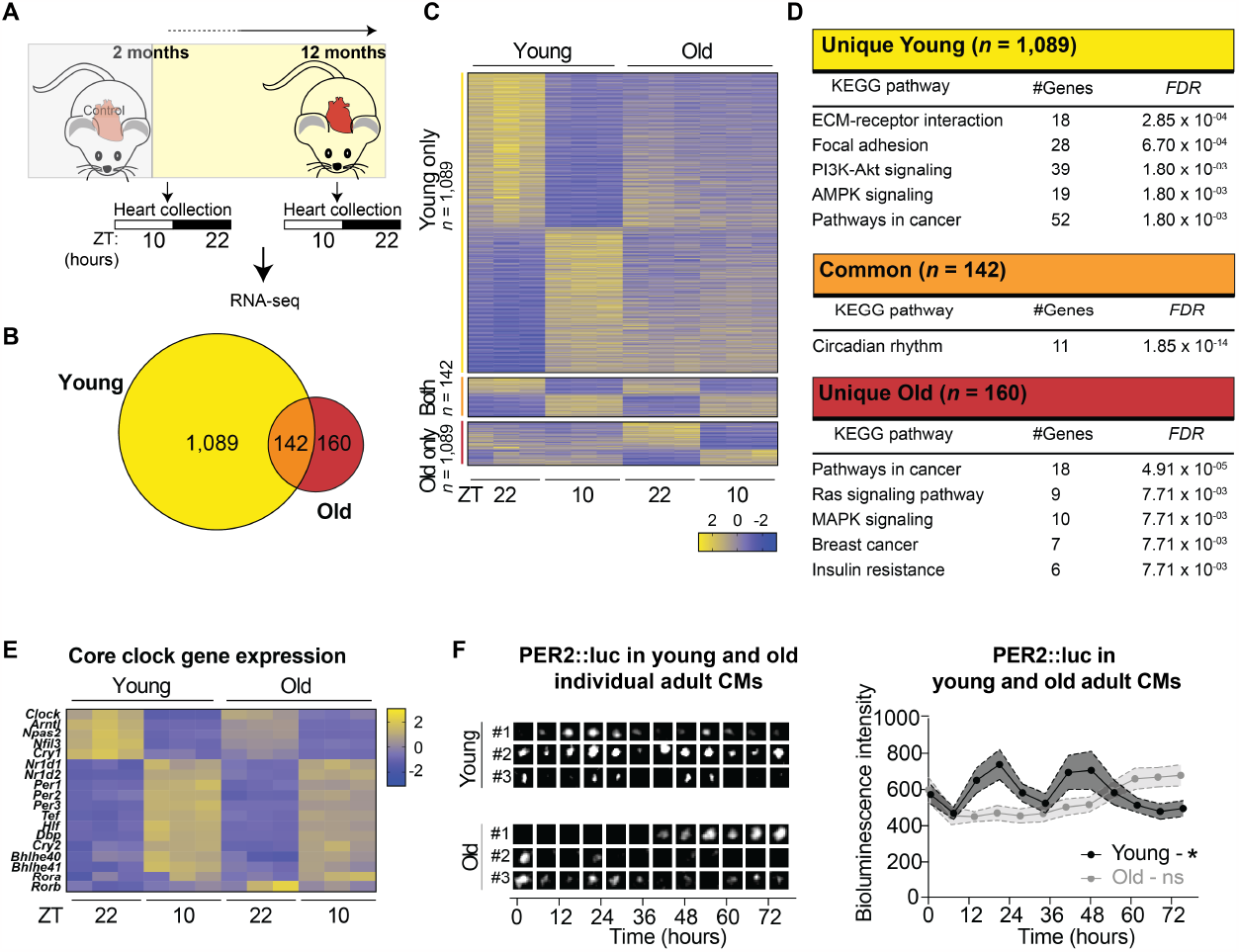
The cardiac diurnal clock is reprogrammed during aging. **a** Schematic representation of experimental setup in female mice. **b** Venn diagram showing the overlap between oscillators in young (2months old, left) and old (1y-old, right) female hearts (*n* = 3/condition, cut-off: DESeq2^40^ Adj.*p* Adj. *P* (FDR) < 0.05) **c** Heatmap depicting z-normalized expression levels for genes in (**a**). **d** KEGG pathway analysis on all DEGs from (**b-c**). The analysis was performed via the use of Enrichr. **e** Heatmap depicting z-normalized expression levels for core clock genes in young (2 months old, left) and old (1 year old, right) female hearts. **f** Bioluminescent tracks from cardiomyocytes isolated from young and old male Per2::luc animals (left) and their quantification (right, *n* = 10 cells/condition). Circadian oscillations were analyzed using the RAIN algorithm^41^, and the significance of rhythmicity across 72 h is indicated. Graphs show mean ± s.e.m., ns = non-significant; *p < 0.05.

Since the core clock did not change significantly at the transcriptional level, we wanted to test whether the observed differences of rhythmicity of clock output genes was regulated by alterations of the core clock at the protein level. Therefore, we used *Per2::luciferase* (PER2::luc) reporter mice^27^ and isolated cardiomyocytes (CMs) from young (3 months) and old (13 months) mice. Using a bioluminescence microscope that can track luciferase signal continuously over multiple consecutive days, we noted that observed circadian oscillations of young PER2::luc CMs were reduced in old CMs (**Fig. 1f**). Therefore, alterations of PER2 protein levels are likely to contribute to altered core clock function in the heart during aging. This is in line with observations in the liver, in which SIRT1, a deacetylase that uses NAD^+^ as a cofactor, was shown to regulate PER2 localization and stability via deacetylating this clock factor^20^. While this suggests a similar mechanism could possibly function in the heart, many regulatory elements of both rhythmicity and NAD^+^ differ between tissues, so further work is needed to confirm this pathway in a cardiac context.

To investigate the potential link between NAD^+^ and reduced core clock function in the heart, we evaluated cardiac NAD^+^ levels during aging. NAD^+^ levels in old (15 months) female mice were lower compared to young (3m) mice (**Fig. 2a)**. We next wanted to test whether boosting NAD^+^ levels can improve diurnal gene expression in the heart. Therefore, we supplemented the drinking water of 2-month-old female mice with nicotinamide riboside (NR) *ad libitum* for a period of 10 months and harvested animals at the age of 1 year (**Fig. 2b)**. Female mice specifically were chosen due to prior evidence that the cardiac response to NR was beneficial in females, at least in a model of circadian disruption and potentially advanced aging^19^. NAD^+^ levels in the liver were drastically increased (**Supplemental Fig. 2**), whereas hearts only showed a modest trending increase (*p* = 0.055, **Fig. 2c**). These findings are in line with previous results^4,28^ and could be due to high cardiac metabolic turnover^29^. Nonetheless, while body weight was unaltered (**Fig. 2d**), we observed that naturally occurring cardiac enlargement was completely abolished in NR-treated mice (**Fig. 2e**). In line with this, the cardiac stress marker *Anp* was significantly reduced in these mice (**Fig. 2f**), showing that the supplementation strategy does positively affect the heart. This highlights the therapeutic potential of NAD^+^ boosting during aging.

**Fig. 2.**
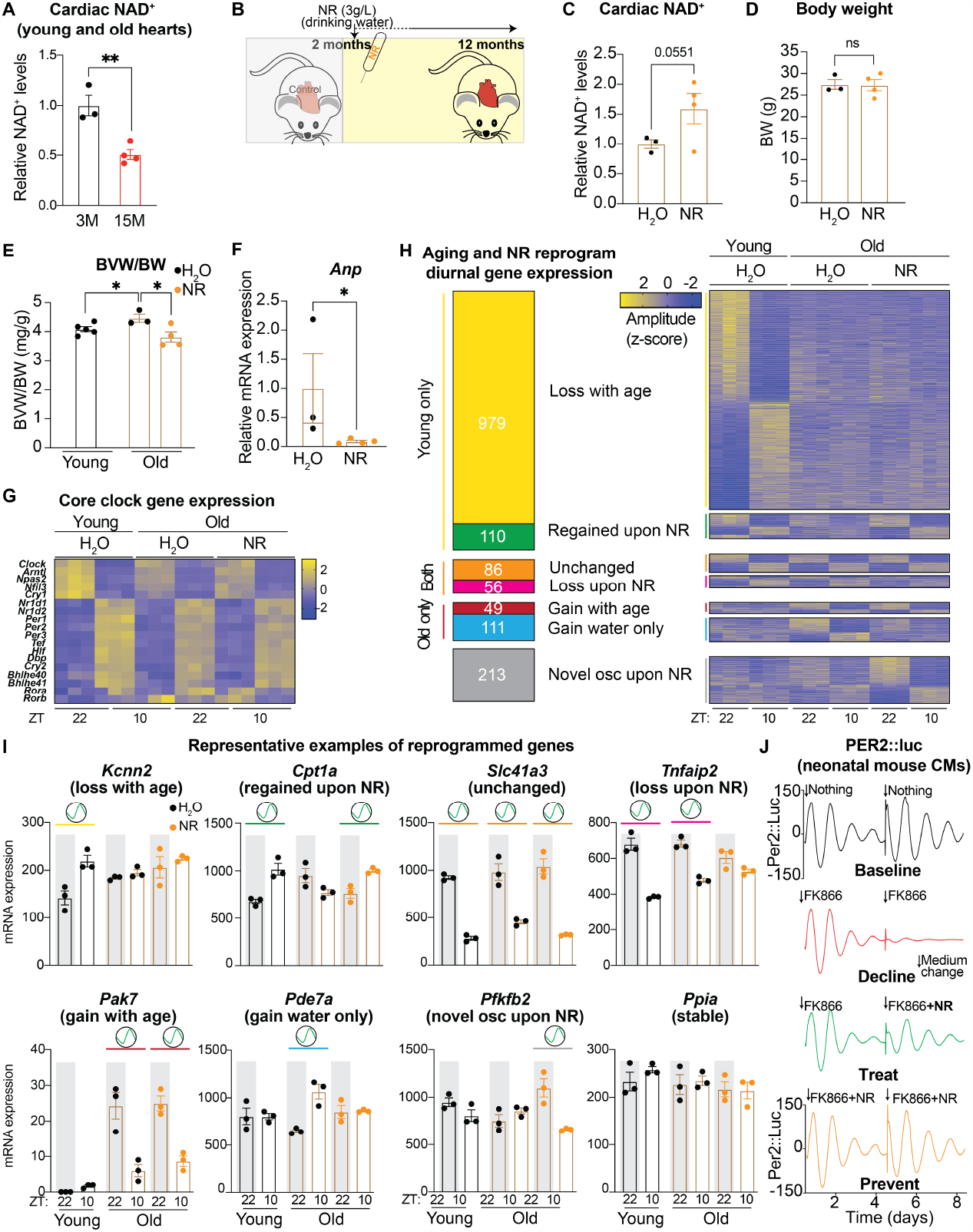
Nicotinamide Riboside supplementation reprograms the cardiac diurnal transcriptome. **a** Cardiac NAD^+^ levels from young (3M) and old (15M) female mouse hearts (*n* = 3-4/condition) at ZT22. **b** Schematic representation of NAD^+^ boosting strategy in female mice with NR. **c** Cardiac NAD^+^ concentration and **d** body weight (BW) of female mice treated with NR versus H2O (control) for 10 months starting at an age of 2 months (*n* = 3-4/treatment) at ZT10. **e** Biventricular to body weight (BVW/BW) ratio in young and old mice treated with NR versus H2O (control) (*n* = 3-5/condition). **f** *Anp* (*Nppa*) mRNA expression in hearts from NR vs control treated mice (*n* = 3-4/condition) at ZT10. **g** Heatmap depicting z-normalized expression levels for core clock genes in young and old NR vs control treated female hearts. **h** Identified categories of oscillators (left) and heat map (right) depicting their z-normalized expression values, depending on their diurnal expression in each condition (ZT10 vs ZT22, DESeq2^40^ Adj.*p* < 0.05, *n* = 3/condition). **i** Gene expression (DESeq2 normalized counts) for representative genes of each class. Graphs show mean ± s.e.m. Oscillation sign depicts differential expression (*Adj. p* <0.05 assessed by DESeq2, *n* = 3/condition). **j** average background-subtracted PER2::luciferase oscillations from neonatal mouse cardiomyocytes treated with NAMPT inhibitor FK866 (100 nM) with or without NR (500 µM). Arrows indicate medium change and addition of fresh FK866 and NR. Cells were derived from *Per2::luciferase* mice and monitored *ex vivo* in a Lumicycle (*n* = 4). *p < 0.05; **p < 0.01; by 2-tailed unpaired Student’s *t*-test.

To investigate whether the clock was affected by NR treatment, we assessed gene expression at ZT22 (night) and ZT10 (day) in old (1 year) NR-treated mice and compared these to young and old untreated mice (**Fig. 1c**). We did not observe major changes in core clock gene expression (**Fig. 2g**). From the genes that maintain rhythmicity upon aging (**Fig. 1c**, *n* = 142), the majority (*n* = 86/142) were unaffected by NR and stayed rhythmic, while 56/142 genes lost rhythmicity (**Fig. 2h-i**). From the genes that lost rhythmicity upon aging (**Fig. 1c**, *n* = 1,089), most were unaltered by NR (*n* = 979), but 110 regained diurnal expression. From the latter 18 out of 110 genes inverted their rhythmic expression pattern. The genes that gained rhythmicity with age (**Fig. 1c**, *n* =160), predominantly lost this pattern upon NR (*n* = 111/160). Next to the previously described genes (**Fig. 1c**), we identified 213 genes that gained *de novo* rhythmicity during aging, specifically upon NR treatment (**Fig. 2h-i**). KEGG pathway and epigenetic Landscape In Silico deletion Analysis (LISA) revealed that genes that lost diurnal expression following NR are enriched for longevity regulating and neurotrophin signaling pathways, with NKX2-5, TBX3, and NR3C1 as putative transcriptional regulators of these genes (**Supplemental Table 1**, *p*<0.01). Transcripts that regained diurnal rhythmicity upon NR encompassed those involved in FoxO, AMPK, and adipocytokine signaling and are potentially regulated by TBX3, NKX2-5, and JARID2. *De novo* oscillators upon NR treatment were primarily involved in Apelin signaling, choline metabolism in cancer, and sphingolipid signaling pathways and are genes that are potentially regulated by TBX3, MEF2A, and SUZ12 (**Supplemental Table 1**). These results suggest that NR reprograms the diurnal transcriptome in the aging heart and that different TFs mediate distinct categories of NAD^+^-mediated reprogramming of the diurnal cardiac transcriptome.

To further investigate the observed improvement of clock function upon NR supplementation, we treated neonatal mouse cardiomyocytes from PER2::luc mice with FK866, a potent inhibitor of NAMPT, the rate limiting enzyme for the production of NAD^+^. Upon FK866, we noted a drastic reduction in PER2::luc rhythms. This reduction in rhythmicity could be completely prevented/reversed by concomitantly supplementing the FK866-treated cells with NR (**Fig. 2j**). In addition, adding NR to cells initially only treated with FK866 could enhance rhythmicity again (**Fig. 2j**), suggesting that NR can be used to treat as well as to prevent reduction of rhythmicity. These results show that PER2 protein rhythms are affected by NAD^+^ in the heart and suggest that NR treatment is able to improve cardiac PER2 rhythms, thereby contributing to transcriptional reprogramming.

In summary, we show that rhythmic clock output gene expression in the heart is reduced and rewired with age and that long-time supplementation with NR can shift gene expression to partially restore a young-like program. While proper amplitudes of circadian rhythms are clearly associated with health^21^, the consequences of rhythmicity in downstream outputs are not obvious in terms of delineating standard physiology, compensation, or pathology. However, in the view that advanced aging is generally a less healthy state, as supported by the association of rhythmic pathways observed in older mice with cancer and insulin resistance, the trend of NR treatment to both maintain young rhythmicity (110 genes) and prevent age-induced rhythmicity (111 genes) is encouraging. Additionally, the genes regaining young rhythmicity correspond to changes in FoxO signaling, a pathway critical to heart development and aging while also possibly interacting with SIRT1^30^. These changes also are seemingly mediated by factors critical to cardiac development like TBX3^31^ and JARID2^32^. Interestingly, the novel oscillators gained on NR treatment were found by LISA to correlate with hypertrophy factors like MEF2A^33^ and SUZ12^34^. Further investigation into these factors and pathways is necessary to identify potential mediators, and assess the tendency of NR treatment to affect gene expression or protein activity against cardiovascular disease. The treatment was safe and effective in female mice and restored naturally occurring cardiac enlargement observed with age, though these effects should continue to be explored across further aging and between genders, as gene expression undergoes altered dynamics throughout these parameters^35^. We have here already demonstrated that: even limited aging alters physiological rhythmic expression and cardiac NAD^+^ levels *in vivo*, which can be mildly reversed by NR treatment; manipulation of cardiomyocyte NAD^+^ is possible via small molecules and affects PER2 rhythmicity; and that cardiomyocyte PER2 levels change over age. While the mechanism in this context remains to be clarified, this provides evidence to support the known but understudied involvement of NAD^+^ in cardiac clock rhythmicity and aging, possibly through a loop such as the NAD^+^-SIRT1-PER2 axis recently discovered in the liver^20^. Future studies will prioritize examination of this particular relationship through protein-level analysis of cardiomyocyte PER2 acetylation in response to aging and NR supplementation. Investigation of additional potential NAD^+^ influences on the cardiac clock such as redox status (by comparing NADH levels) and alternate forms of NAD^+^ consumption (such as PARPs) may reveal cooperative or alternative methods of NAD^+^-mediated rhythmicity, as the regulation of NAD^+^ biosynthesis and the clock are both multivariate and tissue-specific. Indeed, Basse et al. recently demonstrated that the core clock in brown adipose tissue is highly depending on NAD^+^ biosynthesis, while the skeletal muscle clock is largely refractory to NAMPT depletion^36^. Although we here show a clear effect of NAD^+^ modulation on the cardiac clock, to which extent the cardiac circadian clock depends on NAMPT-mediated NAD^+^ biosynthesis needs to be tested. Since NAD^+^ supplementation strategies are regularly proposed to combat aging and age-related diseases (e.g. heart failure), assessing the effects on the circadian clock will be crucial. These findings reveal a clear link between NAD^+^ and cardiac diurnal rhythms and offers potential avenues to treat and prevent cardiovascular diseases appearing with age.

## Abbreviations

Ctrl: control
KO: knock-out
ZT: *Zeitgeber*
NAMPT: nicotinamide phosphoribosyltransferase
Adj.p: Adjusted *p*-value
FDR: false discovery rate
Anp: Atrial natriuretic peptide
Per2: Period2
Kcnn2: Small conductance calcium-activated potassium channel protein 2
Pak7: P21-activated kinase 7
Pfkfb2: 6-phosphofructo-2-kinase/fructose-2,6-biphosphatase 2
Tnfaip2: tumor necrosis factor, alpha-induced protein 2
Slc41a3: solute carrier family 41, member 3
Pde7a: phosphodiesterase 7A
Cpt1a: carnitine palmitoyltransferase 1a
Ppia: peptidylprolyl isomerase A
NAD^+^: nicotinamide adenine dinucleotide
NR: nicotinamide riboside
BVW: biventricular weight
BW: body weight

## Acknowledgements and funding

We kindly thank M.A. Lazar for providing us with mouse material, M. Wiesnet for the help with the Langendorff perfusion, and M.W. Vermunt for carefully and critically reading the manuscript. The authors thank Elysium Health for providing NR at no cost and J. Baur for the advice on the NR experiments. This work was funded by the Deutsches Zentrum für Herz-Kreislauf-Forschung e.V. (DZHK) (K.C, Y.N.M, P.D) and the Cardiopulmonary Institute (CPI).

## Author contributions

P.D. conceived and designed the overall study. P.D., B.J.C., K.C., and S.G. contributed to next-generation sequencing experiments and bioinformatic analyses. Y.N.M. and B.J.C. performed NAD^+^ measurements. P.D., B.J.C., and M.L. performed gene expression measurements, cell culture experiments, animal caretaking, and microscopy experiments.

## Competing interests

All authors declare no competing interests.

**Correspondence and requests for materials** should be addressed to Pieterjan Dierickx.

## Methods

### Animals

Female mice on a mixed C57Bl6J/N strain, representing diversified backgrounds used in previous NR treatments, at the age of 2-12 months were used for all experiments unless stated otherwise in figures and figure legends. They were housed under 12h light/12h dark conditions and fed a standard chow diet *ad libitum* with free access to water. NR (Elysium Health, NY, United States) was added to the drinking water (3 g/L) in light-protected bottles that were changed twice per week, starting at the age of 2 months. Animal experiments for RNA-seq were performed in the lab of Prof. Dr. Lazar. All animal care and use procedures followed the guidelines of the Institutional Animal Care and Use Committee (IACUC) of the University of Pennsylvania and by the responsible Committee for Animal Rights Protection of the State of Hessen (Regierungspraesidium Darmstadt, Wilhelminenstr. 1-3, 64283 Darmstadt, Germany) with project number B2-2018.

### Adult mouse cardiomyocyte isolation

The isolated hearts of male PER2::Luc animals were cannulated via the aorta and rinsed with calcium-free perfusion buffer to remove erythrocytes. The heart was enzymatically digested by perfusion with digestion buffer until it became swollen and turned slightly pale. Atria and outflow tract were removed and the ventricle was dissociated in stop buffer. Isolated myocytes were washed and the Ca^2+^ content was adjusted up to 1 mM stepwise. Cells were centrifuged twice for 1 min at 300 rpm. The cell pellet constituting the cardiomyocyte fraction was taken up in culture medium and seeded on laminin coated dishes. Culture medium was changed after 2–3 hrs. The culture medium of cardiomyocytes was changed after 12–16 h and cells were cultured overnight.

### Microscopic real-time bioluminescence analysis

Bioluminescence from adult PER2::luc cardiomyocytes was assessed with an LV200 microscope (Olympus) in a humidified chamber under 5% CO2, at 37°C guided by a controller (Tokahit), using a 10×0.3NA Plan Semi Apo objective (Olympus). A transmitted light image was recorded prior to the start of each imaging run to localize the cells. Bioluminescence was detected for multiple consecutive days, using an EM-CCD camera (Hamamatsu), with exposure times of 8min, and an interval of 6h. Image series were analyzed in ImageJ. Cells were synchronized with 100 nM dexamethasone for 2h and changed to adult cardiomyocyte culture medium, containing 1mM D-Luciferin Potassium Salt (Promega).

### Neonatal mouse cardiomyocyte isolation

Neonatal mouse cardiomyocytes from P0-P3 *PER2::Luc* animals were harvested with a primary mouse cardiomyocyte isolation kit (Pierce) following the manufacturer’s protocol.

### Bioluminescent recording and data analysis

Neonatal mouse cardiomyocytes were synchronized with 100 nM Dexamethasone (Sigma) for 2 h. Subsequently, medium was changed to recording medium (Powdered DMEM (Corning), 10 mM HEPES, 1% P/S, 5% FBS, 0.035% Sodium bicarbonate, 100 μM D-Luciferin Potassium Salt (Promega) with the addition of 100 nM FK866, FK866 + 500 µM NR, or nothing. Culture dishes were sealed with high vacuum grease (Dow Corning) and monitored via the use of a LumiCycle32 device (Actimetrics) at 37°C. After 4 days, recording medium was refreshed with the addition of fresh FK866, FK866 + NR, or nothing. Bioluminescence from each dish was continuously recorded (integrated signal of 70 s with intervals of 10 min). Raw data (counts/s) were baseline subtracted.

### RNA isolation and quantitative RT–PCR

Total RNA was extracted from heart tissues (TRIZOL) using RNAeasy (Qiagen) according to the manufacturer’s instructions, and treated with DNase (Qiagen) before reverse transcription. cDNA was generated using High-Capacity cDNA Reverse Transcription Kit (Applied Biosystems). Quantitative PCR reactions were performed using PowerSYBR Green PCR Master Mix (Applied Biosystems) with specific primers on a QuantStudio 6 Flex instrument (Applied Biosystems). mRNA expression was normalized to the housekeeping gene *Ppib* for all samples. Primer sequences for qRT–PCR: *Ppib*-fw, 5’-GCAAGTTCCATCGTGTCATCAAG-3’; *Ppib*-rev, 5’-GCTTCCAGGCCATATTGGAGCAAA-3’; *Anp/Nppa-rev*, 5’-TGACCTCATCTTCTACCGGCATCT-3’.

### NAD^+^ measurements

Hearts were harvested and snap frozen in liquid nitrogen. Hearts were powdered and subsequently lysed in 0.6M perchloric acid. Cardiac NAD^+^ levels were measured by a cycling enzymatic assay as previously described^4^. A Tecan Infinite M200 Pro microplate reader with fluorescence excitation at 544 nm and emission at 590 nm was used. NAD^+^ values were normalized to tissue weight.

### RNA sequencing and analysis

RNA and library preparation integrity were verified with LabChip Gx Touch 24 (Perkin Elmer). 1μg of total RNA was used as input for VAHTS Stranded mRNA-seq V6 Library preparation following manufacture’s protocol (Vazyme). Sequencing was performed on NextSeq2000 instrument (Illumina) with 1×72bp single end setup. Trimmomatic version 0.39 was employed to trim reads after a quality drop below a mean of Q15 in a window of 5 nucleotides and keeping only filtered reads longer than 15 nucleotides (Trimmomatic^37^: a flexible trimmer for Illumina sequence data). Reads were aligned versus Ensembl mouse genome version mm10 (Ensembl release 101) with STAR 2.7.10a (STAR^38^: ultrafast universal RNA-seq aligner). Alignments were filtered to remove: duplicates with Picard 3.0.0 (Picard: A set of tools (in Java) for working with next generation sequencing data in the BAM format), multi-mapping, ribosomal, or mitochondrial reads. Gene counts were established with featureCounts 2.0.4 by aggregating reads overlapping exons excluding those overlapping multiple genes (featureCounts^39^: an efficient general-purpose program for assigning sequence reads to genomic features). The raw count matrix was normalized with DESeq2^40^ version 1.36.0 (Moderated estimation of fold change and dispersion for RNA-Seq data with DESeq2). Contrasts were created with DESeq2 based on the raw count matrix. Genes were classified as significantly differentially expressed at average count > 5, multiple testing adjusted p-value < 0.05. The Ensemble annotation was enriched with UniProt data (Activities at the Universal Protein Resource (UniProt)). KEGG pathway enrichment was assessed via Enrichr.

### Statistics

Statistical analyses were performed using Prism (GraphPad Software). All data are reported as mean± s.e.m. A 2-tailed unpaired Student’s *t*-tests was used when comparing two conditions (qRT-PCR and NAD^+^ measurements). Statistical analyses to detect circadian oscillations in PER2::luc levels (LV200) were performed by RAIN^41^. For differential gene expression in the RNA-seq data, *p* values were calculated via DESeq2^40^.

### Data availability

The data that support the findings of this study are available in the main text and mRNA-seq data has been deposited at the NCBI data repository with the BioProject accession number PRJNA1002221:

https://dataview.ncbi.nlm.nih.gov/object/PRJNA1002243?reviewer=321vnp5td9hud78gbitu0r70mp.

## Supplemental Information

**Supplemental Table 1**

**Supplemental Fig. 1.**
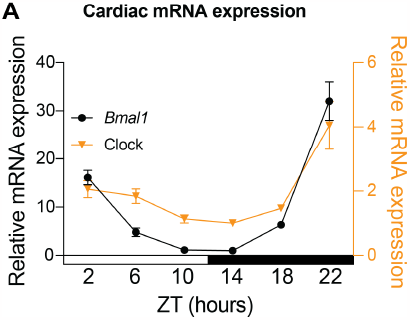
Circadian *Bmal1* and *Clock* mRNA expression in 12-week-old female mouse hearts (*n* = 4/timepoint).

**Supplemental Fig. 2.**
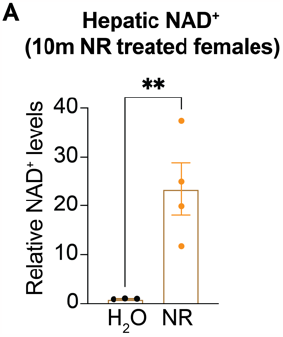
NAD^+^ concentration in livers of female mice treated with NR versus H2O (control) for 10 months starting at an age of 2 months at ZT22 (*n* = 3-4/treatment).

## Notes

### Competing Interest Statement

The authors have declared no competing interest.

